# Targeting neuroinflammation with Abscisic Acid reduces pain sensitivity in females and hyperactivity in males of an ADHD mice model

**DOI:** 10.1101/2022.08.26.505367

**Authors:** Maria Meseguer-Beltrán, Sandra Sánchez-Sarasúa, Marc Landry, Nóra Kerekes, Ana María Sánchez-Pérez

## Abstract

**Aims:** Attention deficit/hyperactivity disorder (ADHD) is a neurodevelopmental syndrome characterized by dopaminergic dysfunction. In this study, we aimed to demonstrate the link between dopaminergic deficit and neuroinflammation underlying ADHD symptoms.

**Subjects and Treatment:** We used a validated ADHD mice model, that involves perinatal 6-OHDA lesion. Animals were treated with 20mg/L (drinking water) of Abscisic acid (ABA) for one month. We tested behaviour (learning and memory, anxiety, social interactions, and pain) in both females and male mice, in all eight groups (control and lesioned, with/without ABA). Postmortem, we analyzed microglia morphology and Ape1 expression in specific brain areas related to the descending pain inhibitory pathway.

**Results:** In females, dopaminergic deficit increased pain sensitivity, but not hyperactivity, in contrast to males. This behaviour was associated with inflammatory microglia and lower Ape1 levels in the anterior cingulate cortex (ACC) and posterior insula cortex (IC). ABA treatment reduced inflammation and alleviated pain. In males, ABA reduced hyperactivity, but had no significant effect on inflammation.

**Conclusions:** This is the first study proving a sex-dependent association between dopamine dysfunction and inflammation in specific brain areas, leading to different behavior outcomes in a mouse model of ADHD. These findings provide new clues for potential treatments.

## 1. Introduction

Attention deficit/hyperactivity disorder (ADHD) is a neurodevelopmental syndrome, affecting 5-10% of children and 2,5% of adults worldwide [1]. ADHD often co-occurs with other psychiatric conditions (i.e. Autism spectrum disorder, anxiety, behavioural problems) [2] and somatic complaints (e.g. asthma [3], celiac disease [4], migraine [5]) which can make diagnosis difficult [6]. ADHD deeply affects patients’ functioning and quality of life (QoL) [7–9]. ADHD’s social impact is also relevant: throughout an individual’s lifetime, ADHD can increase the risk of other psychiatric disorders, educational and occupational failure, accidents, criminality, social disability, and addictions [1,10]. ADHD is usually diagnosed in boys and girls in approximately a ratio of 2:1, with clear phenotypical differences [11]. A recent meta-analysis revealed that girls do not often display the symptoms of hyperactivity and impulsivity, which are more often recognized in boys. However, the executive deficits present in ADHD compared to typically developed controls, are equally distributed amongst girls and boys [12].

The heritability of ADHD is high [13], but environmental factors greatly contribute to its onset and development [14]. Growing evidence points to the influence of inflammation [15,16], microbial dysbiosis [17], and maternal autoimmune reactions [18], affecting ADHD phenotypic variations like anxiety [19], and possibly pain [20]. Systemic inflammation is increasingly acknowledged as a common denominator for psychiatric disorders [21] but the mechanism by which a general inflammatory situation leads to specific psychiatric symptoms remains elusive.

Neuroinflammation is a process underlying brain dysfunction in several neurological and psychiatric disorders [22,23]. Increased inflammation during early neurodevelopment is suspected as a risk factor aggravating and/or triggering ADHD symptoms [24], although no mechanism has been proposed yet. Epidemiological studies including metanalyses reveal that patients with ADHD are more likely to suffer other inflammatory conditions (e.g. asthma, allergic rhinitis, atopic dermatitis, and allergic conjunctivitis) [16,25,26]. Moreover, maternal inflammatory status can prompt the occurrence of ADHD and other neurodevelopmental disorders in the offspring [27,28],[29].

Attention and pain are intimately related [30–32]. In humans, evidence indicates that ADHD increases pain perception in adults [33,34] and the prevalence of generalized pain is higher in ADHD patients (up to 80%) compared to control population (17%) [35,36]. Neuroanatomical studies have shown that attention and pain transmission use same neural networks. One of which, the anterior cingulate cortex (ACC) that receives sensitive information via the thalamus and projects to the insular cortex (IC) has been well-characterized [37,38]. ACC hyperexcitability is associated to higher levels of pain [39], and glia activation can trigger hyperexcitability modulating neuronal channel function [40]. Furthermore, inflammatory markers in ACC are linked to pain sensitization [41–43], thus we aimed to test whether dopaminergic dysfunction underlying ADHD symptoms could cause ACC inflammation, so, consequently reducing inflammation would alleviate ADHD symptomatology. To that end we evaluated microglia morphology and oxidative stress markers in the ACC and insular cortex in relation to behaviour in a previously validated mice model of ADHD [44]. In this model, we tested the effects of ABA alleviating inflammation and behavior. Animal models of ADHD have mostly focused on male subjects. In this study, we studied both females and males, to discern whether the differences in symptoms in girls and boys may have biological grounds, or they are due to gender-stereotypical expressions of common pathophysiology.

## 2. Materials and Methods

### 2.1 Animals and surgical procedures

Female and male Swiss mice (Janvier-Labs; Saint-Berthevin, France) were kept at the animal facility of the University Jaume I. The procedures followed European Community norms on the protection of animals used for scientific purposes. The experiments were approved by the Ethics Committee of the University Jaume I (scientific procedure 2020/VSC/PEA/0099). The animals were maintained on a 12h:12h light cycle and provided with food and water *ad libitum*. Pups were housed with their mothers and kept at constant temperature conditions (24 DC □ 2). After weaning, animals were housed in groups of 2-4 mice to reduce isolation-induced stress. Sixty-nine mice were randomly assigned to sham or 6-OHDA groups. At postnatal day 5 (P5), pups received 6-OHDA hydrobromide (Sigma-Aldrich, France) or vehicle in one of the lateral ventricles as described in [44]. Thirty minutes before surgery, mice were injected with desipramine hydrochloride pretreatment (20 mg/kg sc; Sigma-Aldrich, France), an inhibitor of the noradrenergic reuptake. Mice were anesthetized with 3% isoflurane and maintained with 0.8% isoflurane during surgery. 25 μg of 6-OHDA dissolved in 3 μl ascorbic acid 0.1% or vehicle were infused into one of the lateral ventricles (stereotaxic coordinates: AP −2 mm, ML ±0.6 mm, DV −1.3 mm from bregma) [45]. After weaning (P21), mice were randomly administered either ABA in the drinking water (20mg/L), or vehicle. Animal welfare was monitored all through the procedure, according to Ethical committee rules (timeline Figure 1A).

**Figure 1.**
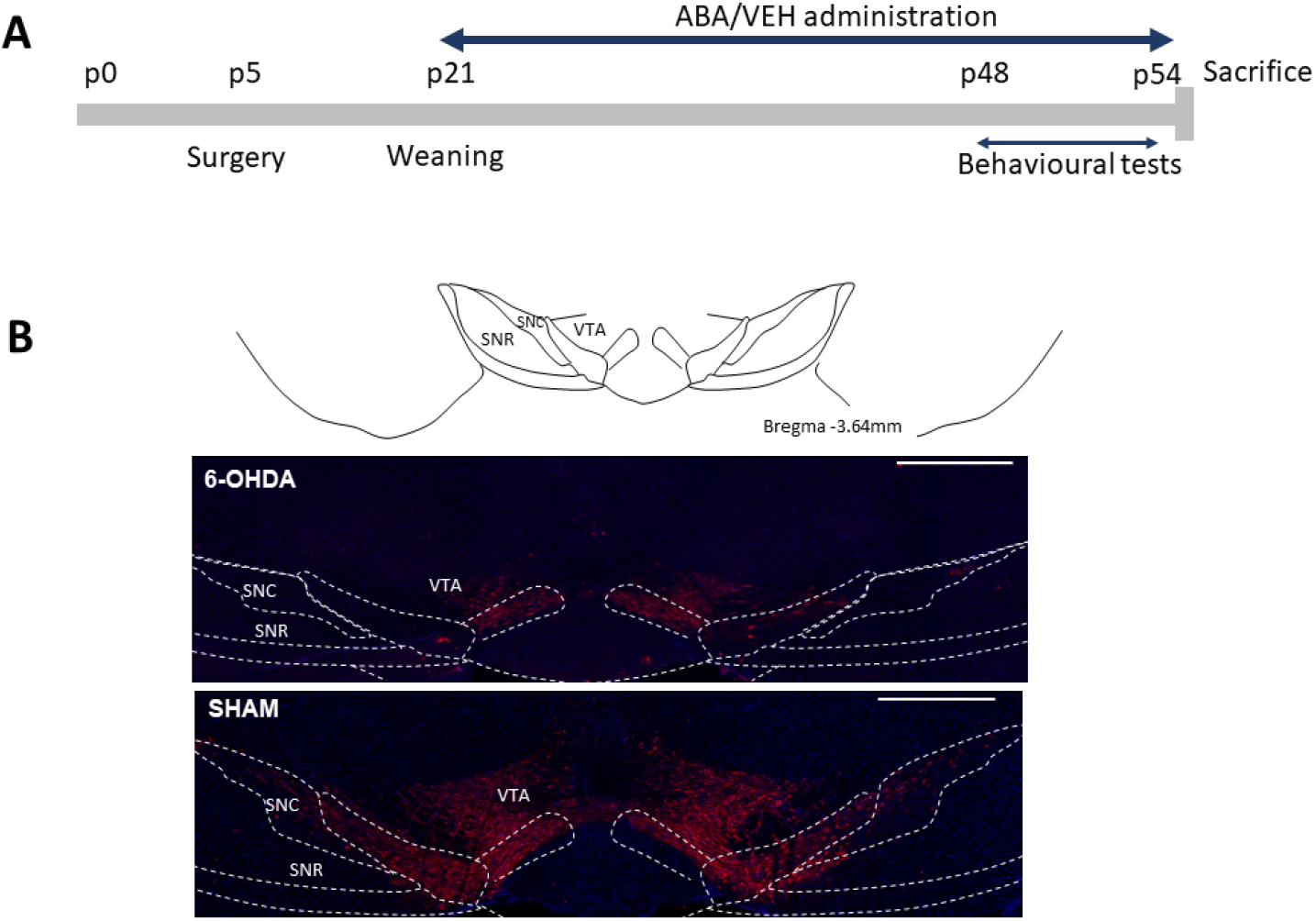
Experiment design. **A)** Timeline of the experiment. Surgery (inoculation of 6-OHDA or sham into lateral ventricle AP −2mm, ± ML 0.6mm, DV −1,3mm from Bregma) was performed on day 5 postnatal (P5). ABA or Vehicle administration started at weaning day, 21 days postnatal (p21) for one month. Behavioural tests were carried out for a week before terminating the experiment. **B)** VTA and SN. VTA, ventral tegmental area; SNC, substantia nigra compact part; SNR, substantia nigra, reticular part. Calibration bar; 500μm.

### 2.2 Behavioral procedures

In all the procedures mice were habituated to the testing room 30 minutes before performing each behaviour paradigm. Animals were recorded using a video tracking system (ANY-maze, Stoelting Europe, Dublin, Ireland). All behavioural procedures were conducted in dim light. Behavioural tests started at P48, one week before completing ABA treatment. The researcher was blind to the group’s condition. All apparatus and objects were cleaned with a 30 % ethanol solution between subjects.

#### Spontaneous locomotor activity

was assessed in the Open Field arena (Ugo Basile, Germonio (VA), Italy). Mice were placed in the open field facing one of the walls and allowed to freely explore for 10 min. Distance traveled (cm) and speed (cm/s) were quantified.

#### Recognition memory

was assessed by novel object recognition (NOR) test. NOR experiment was conducted as described [46]. On the test day, the subjects were left to explore two identical objects for 10 min (familiarization phase). After 30 min, mice were back to the arena and allowed to explore one of the previous objects (familiar) and a novel object for 10 min (test phase). “Exploration” is defined as sniffing or touching the object (head oriented towards it). Data were expressed as Discrimination Index (DI), [(Time Spent on the Novel Object – Time Spent on the Familiar Object)/Total exploration time)].

#### Spontaneous alternation

was measured by the T-maze test as described [47]. Briefly, the animal is placed in the starting arm and left to explore for 5 min, with access to two of the three arms (familiarization phase). The mice were then returned to the home cage for a 30 min inter-trial interval and then placed back in the starting position, but now with free access to all three arms for 5 min (test phase). The arm that was closed during the familiarization phase is considered the “novel” arm, and the arm visited during the familiarization phase is considered the “familiar” arm. Data expressed as DI, [(Time Spent on the Novel Arm – Time Spent on the Familiar Arm)/Total exploration time)].

#### Anxiety

was measured through Elevated Plus Maze (*EPM*) (Ugo Basile, Germonio (VA), Italy). EPM is an elevated non-reflective metal plus with two closed arms (35×15×60cm). A**s** described previously [48], all mice were placed in the center of the maze and were allowed to run freely around the maze for 10 min. Data were expressed as the percentage of time exploring the open arms compared to the total time in the EPM.

#### Sociability

was evaluated in a Three Chamber social interaction paradigm. Animals were placed in the central chamber and allowed to freely explore all chambers for 10 min (habituation phase) Immediately after the habituation phase, we perform the test phase (10 min). The mouse can explore either an object or a conspecific protected by a fence, placed in the lateral chambers. The chamber in which the conspecific or object was placed was balanced. In addition, we used 4 different conspecifics (2 males and 2 females) to be available to balance them. Data were expressed as the percentage of time exploring the conspecific mice compared to the total time exploring. Climbing or running around it was not considered exploration.

#### Pain sensitivity by mechanical stimulus (Von Frey test)

The mechanical nociceptive response was assessed using the Von Frey test (Ugo Basile, Germonio (VA), Italy) as described in [49]. Briefly, mice were placed in testing individual cages with a mesh floor for 30 min (habituation). The withdrawal threshold of the paw was set by calibrated Von Frey filaments applied to the ventral surface of the hind paws. 3-5 measurements (each paw) were registered with an interval of 30 sec. The pain threshold is the grams of the filament at which the mouse withdrew its paw.

#### Pain sensitivity by thermal stimulus (Plantar Test)

The thermal nociceptive response was assessed using the Plantar test (Ugo Basile, Germonio (VA), Italy). Mice were placed in testing cages with a glass pane floor for 30 min (habituation). Using an infrared (IRed) generator, the ventral surface of the left and right hind paws was stimulated with an IRed intensity of 50. The cut-off time for finishing the IRed stimulation was set at 15 seconds. The latency to the paw withdrawal was recorded and 5 measurements per paw were done with an interval of 2 minutes interval.

### 2.4 Immunofluorescence procedure

Immunofluorescence was performed as described [50]. Briefly, mice were anesthetized and perfused with saline (0.9% NaCl) followed by fixative (4% paraformaldehyde in 0.1 M PB, pH 7.4). After perfusion, the brains were removed, postfixed overnight, and cryoprotected in 30% sucrose in 0.01M PBS pH 7.4 for 3 days. The brains were cut in the rostro caudal direction (40 μm) using a sliding microtome Leica SM2010R (Leica Microsystems, Heidelberg, Germany). Primary antibodies mouse anti-Tyrosine Hydroxylase (Sigma-Aldrich, France; 1:5000); rabbit anti-Iba1 (FUJIFILM Wako Chemicals Europe GmbH, Deutschland; 1:1000), and mouse anti-Ref1/Ape1 (Santa Cruz Biotechnology, Santa Cruz, CA, USA; 1:200) were incubated overnight. Next, sections were rinsed and incubated for 2 h at RT with donkey anti-mouse Cy3 or donkey anti-rabbit Alexa 488 secondary antibodies (Jackson Immunoresearch, Suffolk, UK). Finally, sections were mounted on slides and covered using Fluoromount-G mounting medium (Invitrogen, California, USA).

#### Imaging and analysis

Fluorescence images were taken with a confocal scan unit with a module TCS SP8 equipped with argon and helio-neon laser beams attached to a Leica DMi8 inverted microscope (Leica Microsystems). Excitation and emission wavelengths for Cy3 were 433 and 560–618 nm respectively; Alexa488 labeled excitation wavelength was 488 nm and its emission at 510–570 nm. For the quantification of Tyrosine Hydroxylase labeling, we used a 10x lens. Image J software was used to count Tyrosine Hydroxylase labeling. Data were expressed as the percentage of Tyrosine Hydroxylase labeling with respect sham group. For the quantification of Iba1 labeling, we used 20×lens. Image J software combined with FracLac plugin [51] was used to analyse the microglia morphology in sections from sham and 6-OHDA groups of both females and males as previously described [46]. The microglia morphological parameters that were analysed were (1) <u>cell area</u>, meaning the total number of pixels corresponding to the area occupied by the cell, soma, and branches; and (2) <u>cell perimeter</u>, based on the single outline cell shape. For the quantification of Ref1/Ape1 labelling, we used a 20x lens. Data were expressed as the mean of grey values per area (insula and ACC).

### 2.5 Statistical analysis

The analysis was carried out with Graph Pad software (GraphPad Prism V8 software, GraphPad, La Jolla, CA, USA). Data were subjected to the Shapiro-Wilk test for Gaussian distribution. If normality was confirmed, data was reported as the mean±SEM and the “n” the number of independent subjects. For differences either a 2-WAY ANOVA was carried out to evaluate the interaction between lesion and treatment, and/or a two-tailed Student’s t-test with the probability set at α<0.05 was used.

## 3. Results

### 3.1. 6-OHDA reduced TH staining in the ventral tegmental area

To ascertain the level of dopaminergic lesion induced by 6-OHDA injection, we carried out a TH immunodetection (Figure 1B). We confirmed that in sham surgery animals, the TH expression was detected in the ventral tegmental area (VTA) and adjacent substantia nigra (SN). Animals with less than 45% lesion were discarded from the analysis.

### 3.2. Effect of perinatal icv 6-OHDA injection and ABA treatment on behaviour

#### 6-OHDA lesion increased spontaneous activity in males, not in females. ABA treatment only affected lesion mice in a sex-dependent manner

In females (Figure 2A) two-way ANOVA analysis revealed that there was an interaction between lesion and treatment, (F_(1,36)_= 6.42, p=0.0158). Tukey’s multiple comparisons test revealed that lesion only affected velocity in ABA-treated mice [6-OHDA-ABA vs. SHAM-ABA: q=6.057, p=0.0007,***] and ABA treatment increased velocity only in lesioned females [6-OHDA-VEH vs. 6-OHDA-ABA: q=4.456, p=0.0165, *]. Similarly, distance traveled (Figure 2B) was affected by the lesion (F_(1,36)_=6.86, p=0.0128) only in ABA-treated mice [6-OHDA-ABA vs. SHAM-ABA: q=6.015, p=0.0008,***], and ABA increased distance travelled only in lesioned mice [6-OHDA-VEH vs. 6-OHDA-ABA: q=4.475, p=0.0159, *]. However, in males (Figure 2C), no interaction between lesion and ABA was found. Lesion affected velocity (F_(1,22)_= 5.813, p=0.0247); but ABA reduced it F_(1,22)_= 7.986, p=0.0098. Similarly, distance travelled (Figure 2D) was increased by lesion (F_(1,22_ = 6.129, p=0.0215); and ABA treatment reduced it F_(1,22)_= 8.378, p=0.0084). Tukey’s multiple comparisons test confirmed that 6-OHDA increased spontaneous activity in the untreated group, velocity [6-OHDA-VEH vs. SHAM-VEH: q=4.061, p=0.0409, *] and distance travelled [6-OHDA-VEH vs. SHAM-VEH: q=4.171, p=0.0347, *]. Whereas in ABA-treated male mice, the lesion did not induce a significant increase in velocity [6-OHDA-ABA vs. SHAM-ABA: q=0.7608, p=0.9488, ns] nor in distance travelled [6-OHDA-ABA vs. SHAM-ABA: q=0.7801, p=0.9451, ns]. Finally, the effect of ABA treatment was compared in 6-OHDA males and a reduction in velocity [6-OHDA-VEH vs. 6-OHDA-ABA: q=4.659, p=0.0162, *] and distance travelled [6-OHDA-VEH vs. 6-OHDA-ABA: q=4.778, p=0.0134, *] was observed.

**Figure 2.**
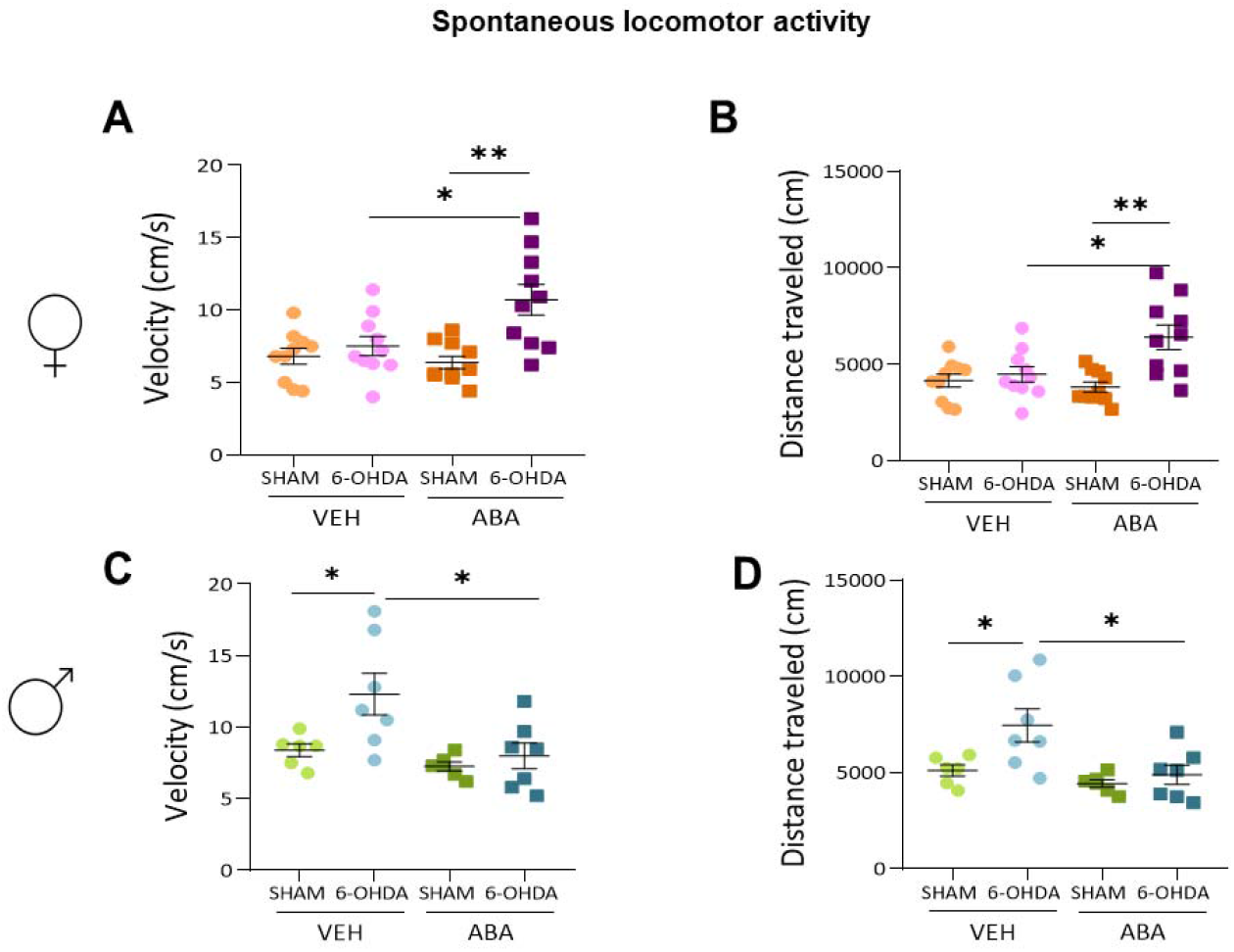
Spontaneous locomotor activity. **A.** Speed (cm/s) and **B.** distance (cm) in females. **C.** Speed (cm/s) and **D.** distance (cm) in males. Data presented as mean ± SEM (n = 6–11 per condition) and analyzed by 2way ANOVA followed by Tukey multiple comparisons test (ns, non significant; *p < 0.05; **p < 0.01).

#### 6-OHDA lesion reduces Novel Object Recognition (NOR) but ABA treatment does not rescue the lesion effect

In females (Figure 3A) lesion affected NOR (F_(1,37)_ = 14.94, p=0.0004), whereas ABA treatment did not rescue it (F_(1,37)_= 0.6095, p=0.4400). There were no interaction between lesion and treatment (F_(1,37)_= 0.02032, p=0.8874). Tukey’s test multiple comparisons revealed that the lesion effect was significant in ABA-treated mice [6-OHDA-ABA vs. SHAM-ABA: q=4.055, p=0.0329, *]. We found a statistical difference by Student t-test between untreated sham and 6-OHDA (p=0.023, t=2.465 df=20). Similar data was obtained with males (Figure 3B), lesion reduced the exploration time of the novel object (F_(1,24)_= 16.98, p=0.0004), but ABA treatment had no effect (F_(1,24)_= 1.786, p=0.1940); and there were no interaction between these two factors (F_(1,24)_= 0.05701, p=0.8133). Multiple comparisons Tukey test indicated the lesion had effect only in untreated mice [6-OHDA-VEH vs. SHAM-VEH: q=4.483, p=0.0201, *] not in ABA treated mice and [6-OHDA-ABA vs. SHAM-ABA: q=3.781, p=0.0598, ns]). The difference between ABA sham and ABA lesion was significant by Student t-test (P=0.011, t=3,054, df=11).

**Figure 3.**
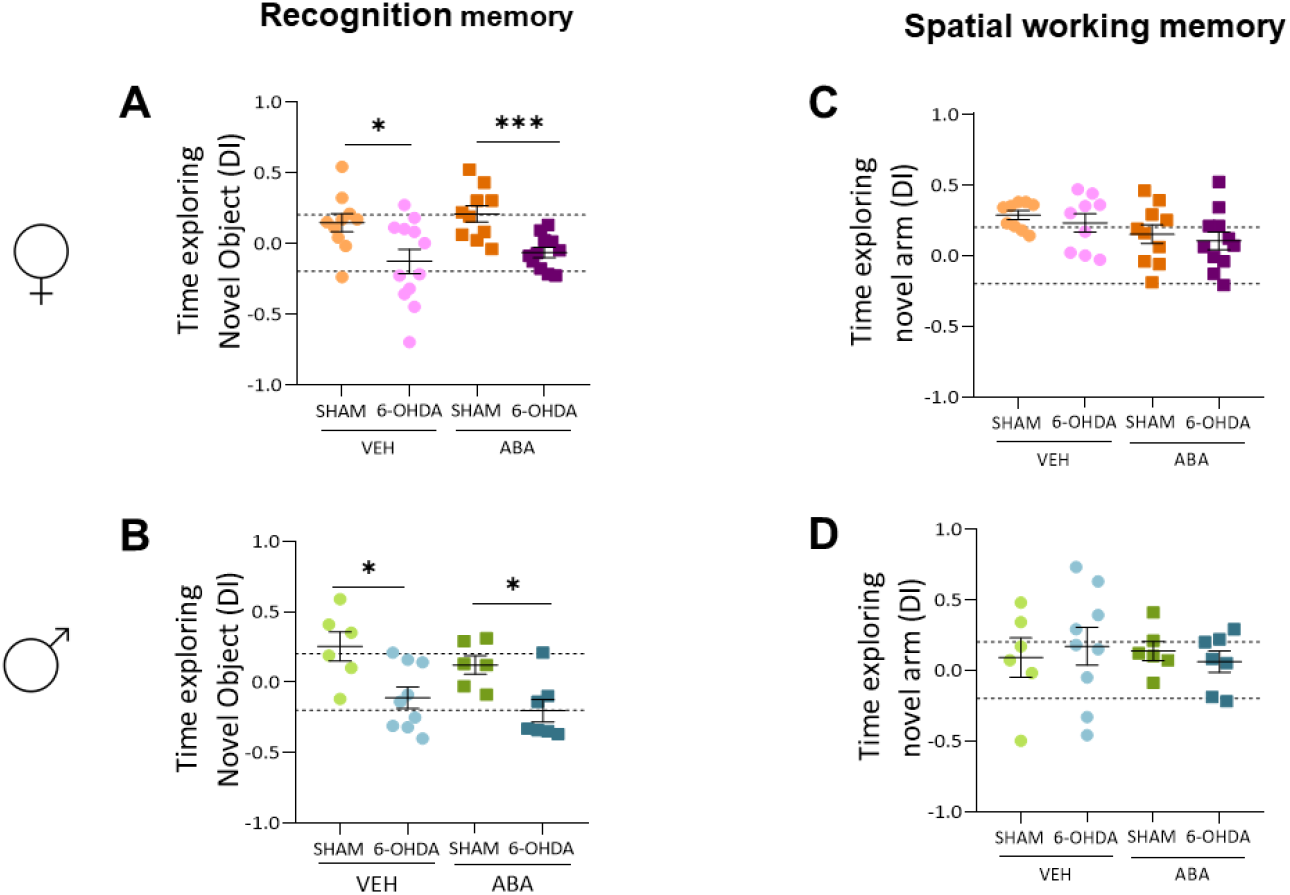
Recognition and spatial working memory. **A)** Time exploring (d-index) the novel object in the Novel Object Recognition test in females and **B)** males. **C)** Time (d-index) exploring the novel arm in the T-maze test in females and **D)** males. Data are expressed as discrimination index [(Time exploring novel – time exploring familiar)/total time exploring], presented as mean ± SEM (n = 6-11 per condition) and analyzed using 2way ANOVA and Tukey’s test, or by two tail Student t-test where indicated (*p < 0.05; ***p < 0.001).

#### 6-OHDA lesion does not alter spatial memory as measured by Spontaneous alternation (T-maze) in females or males. ABA had a sex-dependent effect

In females (Figure 3C) lesion did not affect spontaneous alternation in females (F_(1,35)_= 0.7268, p=0.3997); and surprisingly, ABA treatment had an overall effect (F_(1,35)_= 4.814, p=0.0350). There was no interaction between lesions and ABA treatment (F_(1,35)_= 0.003515, p=0.9531), and Tukey’s multiple comparison test did not show specific ABA effect in Sham, nor lesioned females, suggesting an overall effect. On the other hand, in males (Figure 3D), nor lesion (F_(1,24)_= 0.0004132, p=0.9839), nor ABA treatment (F_(1,24)_= 0.06984, p=0.7938) had effect on spontaneous alternation, suggesting a sex-dependent effect of ABA.

#### 6-OHDA lesion does not alter anxiety as measured by Elevated Plus Maze (EPM) in females or males. ABA displays a sex-dependent effect

Anxiety was not affected in females (Figure 4A) by lesion (F_(1,36)_= 0.2195, p=0.6423), nor treatment (F_(1,36)_= 4.047, p=0.0518). Similarly, lesion (F_(1,24)_= 0.5399, p=0.4696) or treatment (F_(1,24)_= 0.05580, p=0.8153) did not affect anxiety in males (Figure 4B).

**Figure 4.**
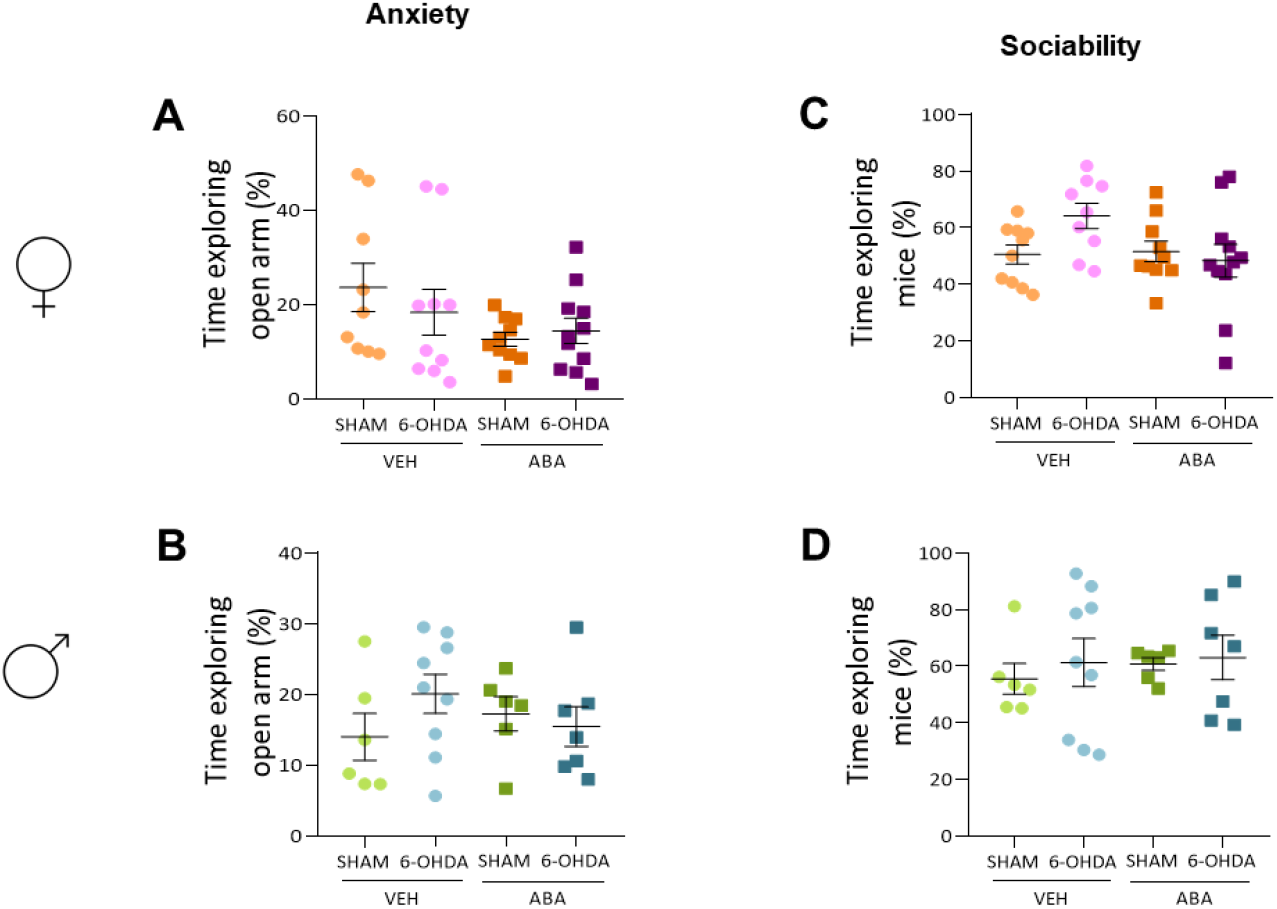
Anxiety and sociability. **A)** Percentage of time to the open arms in the Elevated Plus Maze test in females and **B)** males. **C)** Percentage of time exploring the co-specific mice in the Three Chamber test in females and **D)** males. Data presented as mean ± SEM (n = 6–11 per condition) and analyzed using 2way ANOVA and Tukey’s test. ns, non significant; *p < 0.05.

#### 6-OHDA lesion nor ABA treatment affected Sociability as measured by Three Chamber

In females (Figure 4C) no effect on sociability was found by 6-OHDA lesion (F_(1,36)_= 1.331, p=0.2563), nor ABA treatment (F_(1,36)_= 2.699, p=0.1091). And there was no interaction between these two factors (F_(1,36)_= 3.508, p=0.0692). Similarly, in male (Figure 4D) we observed no effect of lesion (F_(1,24)_= 0.3014, p=0.5881), ABA treatment (F_(1,24)_= 0.2295, p=0.6363). No interaction was found between these two factors (F_(1,24)_= 0.05641, p=0.8143).

#### 6-OHDA lesion had a sex-dependent effect on pain sensitivity by the mechanical stimulus as measured by the Von Frey test

In females (Figure 5A) 6-OHDA lesion increased mechanical sensibility (F_(1,42)_= 4.974, p=0.0311). ABA treatment did not have an overall effect (F_(1,42)_= 0.1429, p=0.7073), and no interaction was found (F_(1,42)_= 1.463, p=0.2332). Tukey’s multiple comparison tests showed no specific differences. However, Student t-test analysis revealed a significant reduction in pain threshold only between vehicle groups (p=0.025, t=2,424, df=20), but not when administered ABA, suggesting that ABA could counteract the 6-OHDA lesion effect.

**Figure 5.**
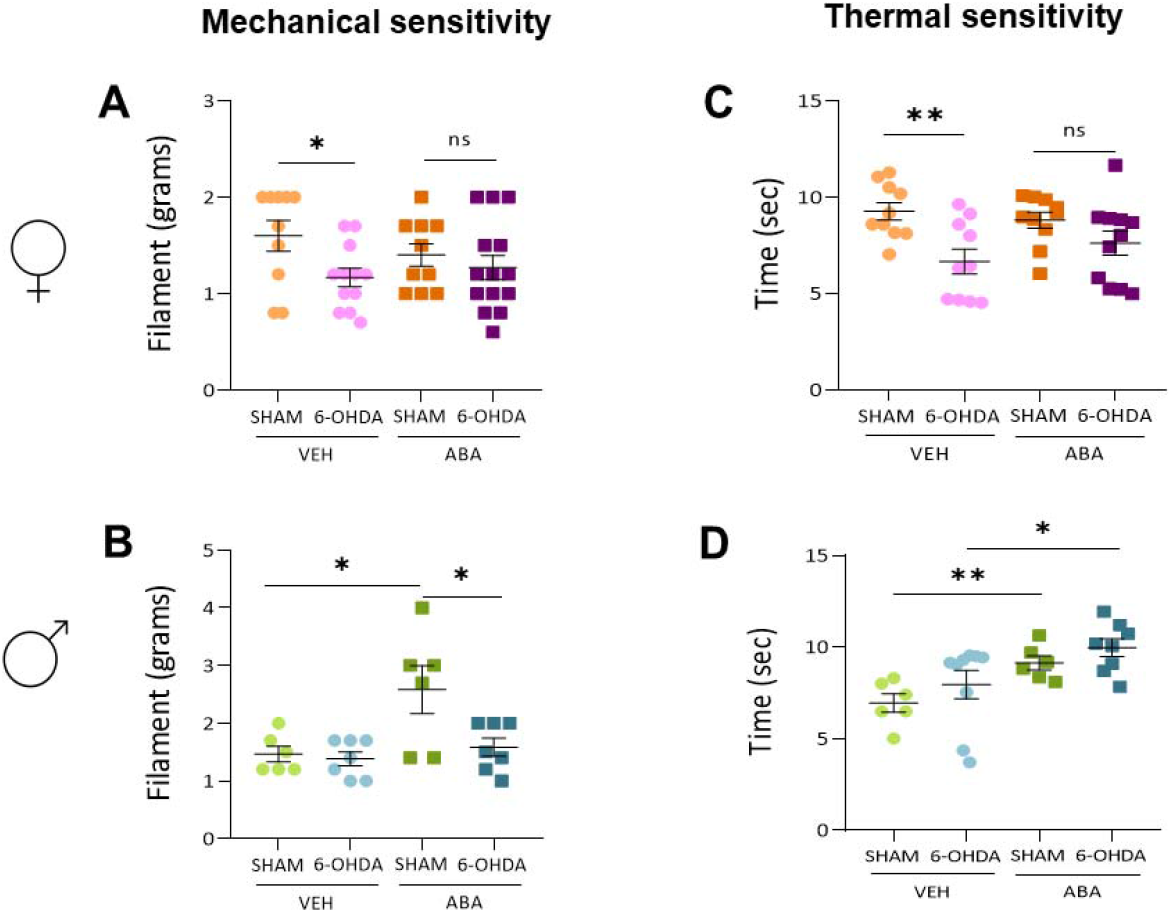
Thermal and mechanical sensitivity. **A)** Threshold (grams) after application of mechanical stimulus in Von Frey test in females and **B)** males. **C)** Threshold (sec) after application of thermal stimulus in Plantar Test in females and **D)** males. Data presented as mean ± SEM (n - 6-11 per condition) and analyzed using 2way ANOVA and Tukey’s test. Or Student t-test (ns, non-significant; *p < 0.05; **p < 0.01).

In males (Figure 5B), as in females, no interaction between lesion and ABA was detected (F_(1,22)_= 4.036, p=0.0570). Tukey’s multiple comparison test revealed that ABA increased pain threshold only in Sham mice [SHAM-VEH vs. SHAM-ABA: q=4.717, p=0.0147], and [6-OHDA-ABA vs. SHAM-ABA: q=4.373, p=0.0254,*], suggesting that lesion with 6-OHDA can prevent ABA effect.

#### 6-OHDA lesion had a sex-dependent effect on pain sensitivity by thermal stimulus measured by Plantar Test. ABA treatment reduces sensitivity in both females and males

In females (Figure 5C) lesion decreased pain threshold (F_(1,37)_= 12.29, p=0.0012); however ABA treatment did not (F_(1,37)_= 0.1738, p=0.6792). We did not find interaction (F_(1,37)_= 1.755, p=0.1934). Tukey’s multiple comparisons test showed that lesion reduced pain only in untreated female only [6-OHDA-VEH vs. SHAM-VEH: q=4.775, p=0.0090, **], suggesting that ABA treatment prevented lesion effect.

On the contrary, in males (Figure 5D), 2-way ANOVA showed that lesion had no effect (F_(1,25)_= 2.189, p=0.1515), but ABA reduced sensitivity (F_(1,25)_= 11.43, p=0.0024). No interaction between these two factors was observed (F_(1,25)_= 0.02371, p=0.8789). Two-tail Student t-test revealed that ABA improves sensitivity in sham (p=0.0058, t=3.495, df=10) and also in 6-OHDA lesioned (P=0.0473, t=2.16, df=15).

### 3.3. Effect of perinatal icv 6-OHDA injection and ABA treatment on neuroinflammation and oxidative stress, in specific brain areas

#### 6-OHDA lesion induces microglia change to a proinflammatory status in the anterior cingulate cortex of both females and males. ABA rescues the effect of 6-OHDA lesion in females, but not in males

Microglia were visualized by Iba-1 staining in the anterior cingulate cortex (ACC) of females and males (Figure 6A). Quantification a statistical analysis revealed that in females (Figure 6B) 6-OHDA lesion reduced perimeter (Fig. 6B; F_(1,37)_= 4.808, p=0.0347) and area (Fig. 6C; F_(1,37)_= 7.333, p=0.0102) in ACC. Indicating a proinflammatory status induced by the lesion. Tukey’s multiple comparisons test revealed this effect occurs only in untreated females’ perimeter (Fig. 6B [6-OHDA-VEH vs. SHAM-VEH: q=4.335, p=0.0202, *]) and area (Fig. 6C [6-OHDA-VEH vs. SHAM-VEH: q=4.458, p=0.0162, *]) suggesting that ABA prevents lesion proinflammatory effect. Moreover, Two tail Student t-test indicates that ABA significantly improved microglia perimeter (p=0.0343, t=2.281, df=19). In males’, 2-way ANOVA revealed that lesion reduces perimeter (Fig. 6C; F (1, 24) = 17,05P=0,0004) and area (Fig. 6E; F (1, 24) = 11,49P=0,0024). There was no interaction between 6-OHDA and ABA in either perimeter (F_(1,24)_= 0.03088, p=0.8620) not area (F_(1,24)_= 0.001076, p=0.974).

**Figure 6.**
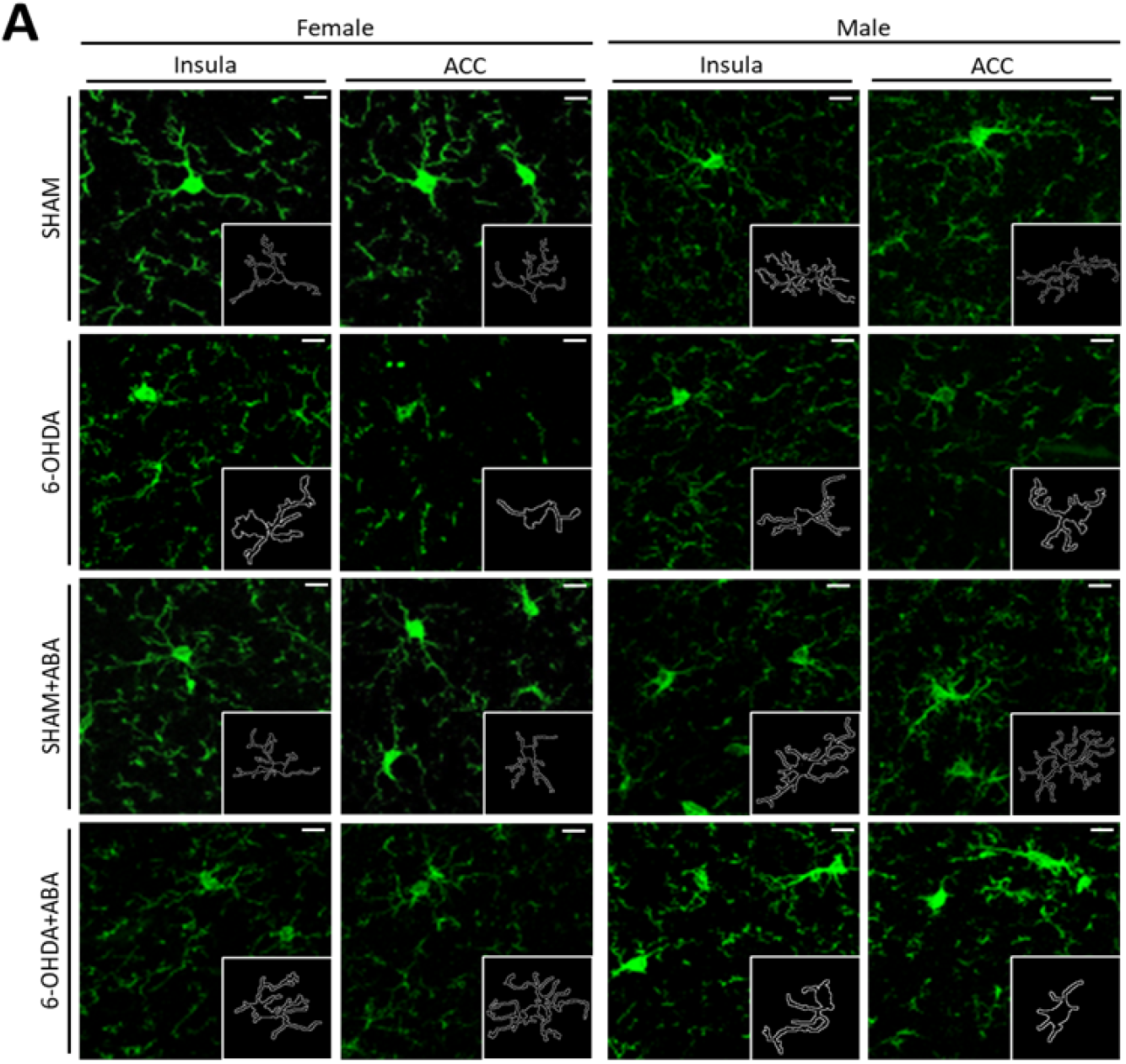
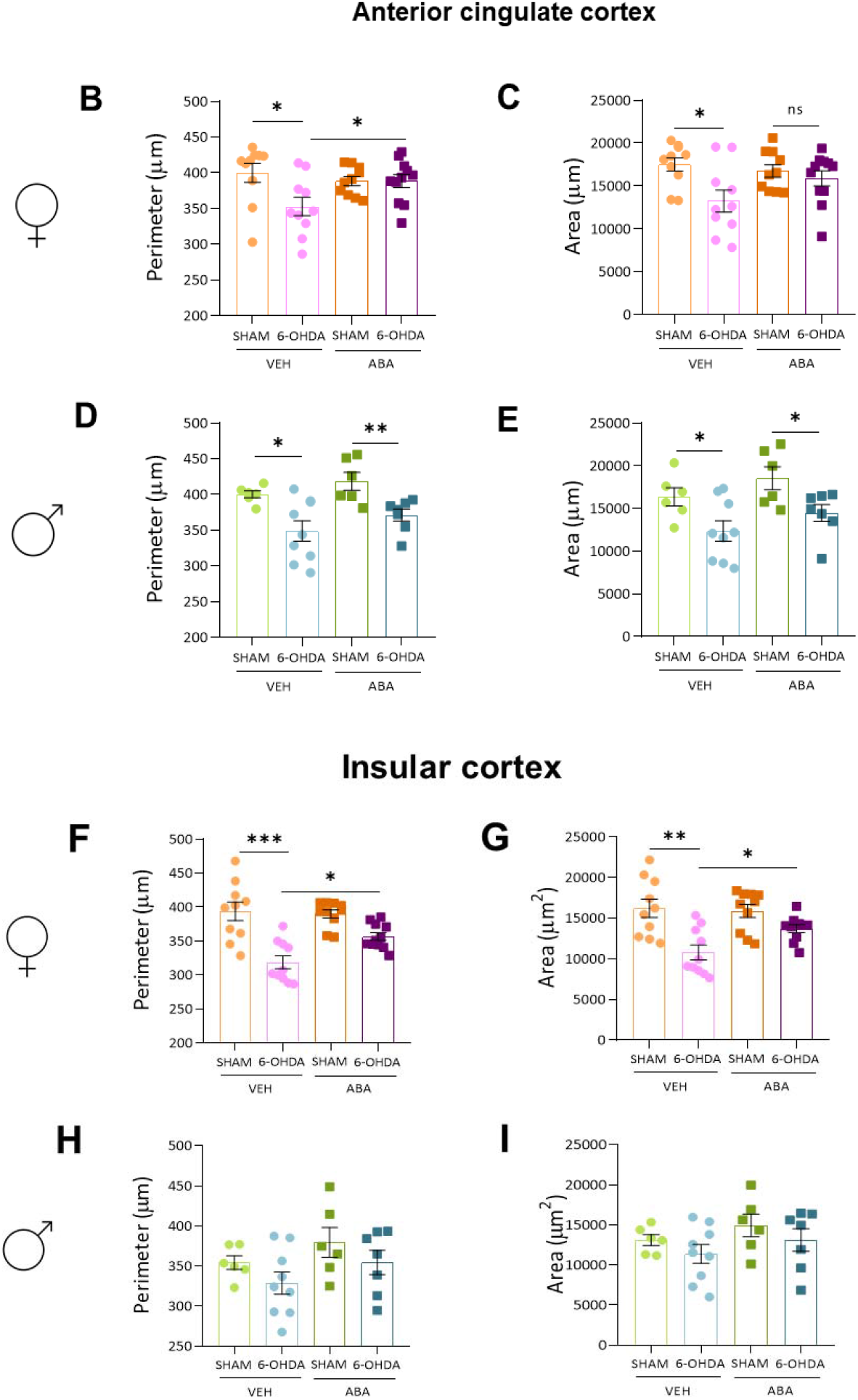
Microglia morphology in Anterior cingulate cortex (ACC) and posterior insular cortex (pIC). **A)** Representative image of microglia in ACC and pIC. **B)** Perimeter (μm) and **C)** area (μm^2^) in ACC in females. **D)** Perimeter (μm) and **E)** Area (μm^2^) in ACC in males. **F)** Perimeter (μm), **G)** Area (μm^2^) in females’ pIC; **H)** Perimeter (μm) **I)** Area (μm^2^) in males’ pIC. Data are expressed as mean ± SEM (n = 6–11 per condition) and analyzed using 2way ANOVA and Tukey’s test, or Student t-test (ns, non significant; *p < 0.05; **p < 0.01; ***p < 0.001). Calibration bar; 10μm.

Two tail Student t-test showed significant differences in the perimeter caused by lesion, in both VEH (p=0.0143, t=2.343, df=13) and ABA treated (p=0.0087, t=3.183, df=11). Similarly, in the area two tail Student t-test showed significant differences in both VEH (p=0.0357, t=2.825, df=13) and ABA treated (p=0.0303, t=2.486, df=11); indicating that ABA could not counteract proinflammatory microglia status induced by 6-OHDA lesion in male’s ACC.

#### 6-OHDA lesion induces microglia change to a proinflammatory status in the posterior insular cortex only in females. ABA rescues the effect of 6-OHDA lesion

In females’ posterior insula 6-OHDA lesion decreased the microglia cells perimeter (Figure 6F) F_(1,37)_= 35.21, p<0.0001. Lesion and ABA treatment had a significant interaction F_(1,37)_= 5.216, p=0.0282). Tukey’s multiple comparisons test revealed that lesion affects microglia morphology in untreated females [6-OHDA-VEH vs. SHAM-VEH: q=8.124, p<0.0001, ****]. Moreover, ABA rescued microglia in 6-OHDA [6-OHDA-VEH vs. 6-OHDA-ABA: q=4.204, p=0.0255, *]. Similar results were observed for the area parameter (Figure 6H). 2-way ANALYSIS revealed lesion effect (F_(1,36)_= 19.83, p<0.0001). Tukey’s multiple comparison test confirmed that lesion effect only on untreated mice [6-OHDA-VEH vs. SHAM-VEH: q=6.374, p=0.0004, ***]. Furthermore, ABA rescued microglia, since a significant difference between lesion-ABA treated and lesion-VEH was found (2tail-Student t-test, p=0.015, t=2.856, df=18). Altogether, this result confirms that lesion induces inflammation and ABA rescues lesion effect in female brain areas (ACC and posterior IC).

On the contrary, in males, we found no significant effect neither lesion or ABA treatment in perimeter (Figure 6H), nor area (Figure 6I) as shown by 2-way analysis F_(1,24)_= 2.892, p=0.1020 and F_(1,24)_= 2.939, p=0.0994; Fig. 6I; F_(1,24)_= 2.023, p=0.1678 and F_(1,24)_= 2.029, p=0.1672, respectively.

#### 6-OHDA lesion and ABA treatment affected differently Ape1 intensity in the anterior cingulate cortex of females and males

Oxidative stress was evaluated by Ape1 immunostaining in the ACC and posterior IC of both female and male mice (Figure 7). Two-way ANOVA analysis showed that in females (Figure 7B), lesion and ABA treatment did not interact, and lesion affected Ape1 intensity significantly (F_(1,36)_= 6.240, p=0.0172), but the Tukey test did not reveal further differences. However, student t-test analysis uncovered that lesion had a significant effect only in VEH (p=0.00.9, t=3.313, df=18), but not in ABA-treated female mice (p=0.46, t=0.748, df=19), suggesting a potential benefit of ABA administration. On the other hand, in males (Figure 7C), there was a strong interaction between the lesion and ABA (F_(1,24)_= 14.98, p=0.0007), Tukey test revealed a significant effect only in ABA-treated mice [6-OHDA-ABA vs. SHAM-ABA q=9.795, p<0.0001, ****]). Student t-test analysis uncovered that ABA could significantly increase Ape1 in Sham (p=0.0023, t=4.069, df=10), but reduce it in lesion (p=0.0258, t=2,494, df=14). This indicates that ABA could rescue lesion-induced Ape1 decrease in female ACC. However, a completely different effect was observed in males, where ABA can increase Ape1 in Sham, but reduce it in 6-OHDA.

**Figure 7.**
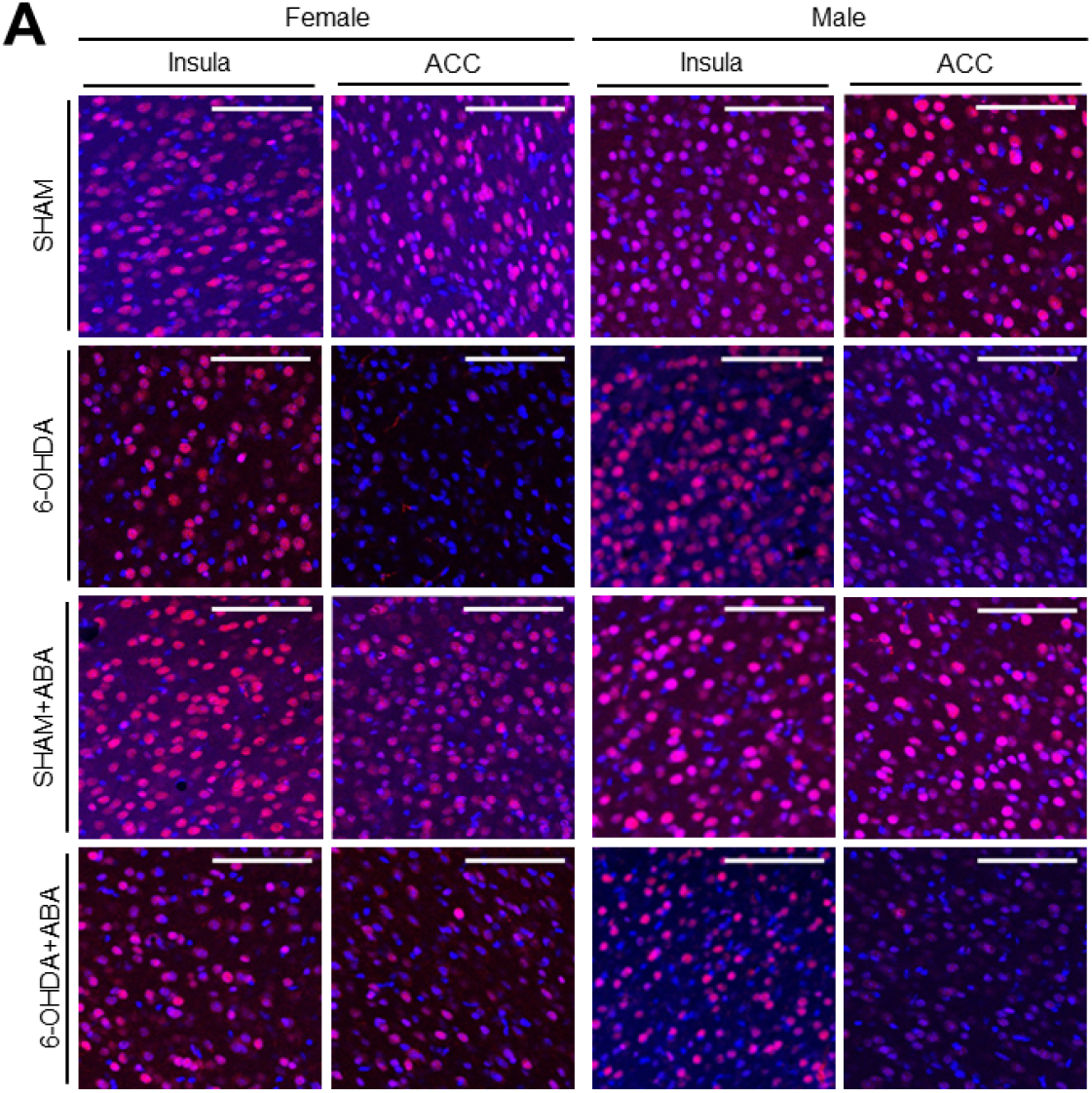
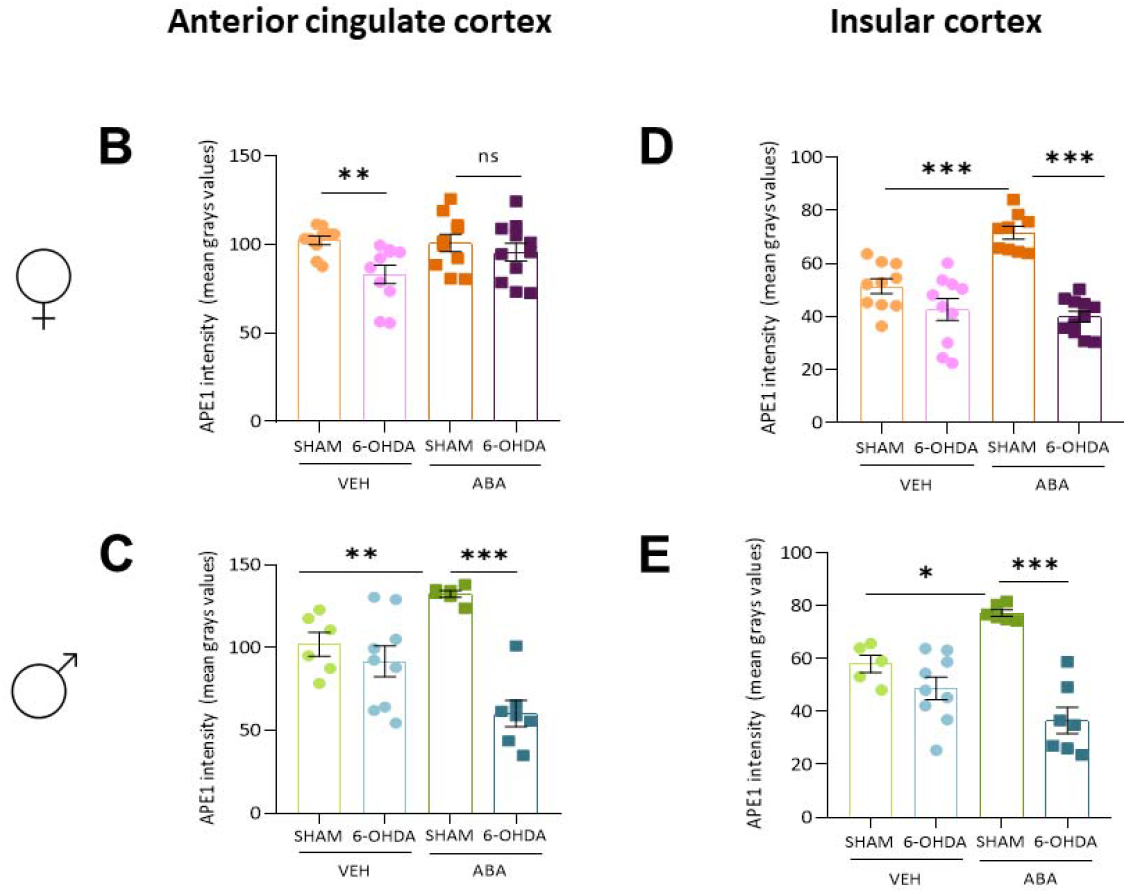
Oxidative stress is measured by Ape1/Ref1 expression. **A)** Representative image of Ape1/Ref1 in the ACC and posterior insula cortex in females and males. **B)** Ape1 intensity in the ACC of females and **C)** males. **D)** APE1 intensity in the insula cortex of females and **E)** males. Data are expressed as mean ± SEM (n = 6–11 per condition) and analyzed using 2way ANOVA and a two-tailed Student t-test. ns, non significant; *p < 0.05; **p < 0.01; ***p < 0.001.

#### 6-OHDA lesion prevents ABA effect on Ape1 intensity in both females’ and males’ posterior insula

When posterior insula was analyzed by two-way ANOVA analysis, we found that in females (Figure 7D) there was a strong interaction between 6-OHDA lesion and ABA administration (F_(1,36)_= 15.33, p=0.0004). Tukey’s multiple comparisons analysis revealed that lesion had a significant effect only in ABA mice [6-OHDA-ABA vs. SHAM-ABA q=10.80, p<0.0001, ****]). Furthermore, ABA increased Ape1 intensity only in Sham [VEH-SHAM vs ABA-SHAM, q=6.735, p=0.0002 ***], and had no effect in lesion females. Similarly, in males’ insula (Figure 7E), there was an interaction between the lesion and ABA treatment (F_(1,23)_= 13.99, p=0.0011). Tukey’s multiple comparisons analysis revealed that lesion had a significant effect only in ABA mice [6-OHDA-ABA vs. SHAM-ABA q=9.723, p<0.0001, ****]) and that ABA only increased Ape1 in Sham mice [VEH-SHAM vs ABA-SHAM q=4.228, p<0.0309, *]. This result suggests that ABA can increase Ape1 intensity in the insula, and the lesion can prevent this ABA effect.

## 4. Discussion

ADHD has been associated with dopamine (DA) system dysfunction for a long time [52,53]. Interestingly, dopaminergic alterations can increase inflammatory conditions [54]. Not surprisingly, ADHD has been associated with inflammatory disorders [16,24], and patients with ADHD often suffer from comorbidities of inflammatory etiology [25,26]. DA synthesis and metabolism not only occur in neurons, but also microglia, astrocytes, and oligodendrocytes. Moreover, these non-neuronal cells express functional D1 and D2 receptors (for review see [55]).

In our study, we tested the hypothesis that targeting neuroinflammation would reduce some ADHD core symptoms, e.g. spontaneous locomotor activity, and comorbid symptoms like anxiety, sociability, and pain sensitivity [20].

We made use of a mouse model of ADHD obtained with perinatal icv injection of 6-OHDA [44] and evaluated whether dopaminergic lesion-induced behavioural alterations were associated with neuroinflammation and oxidative stress in specific brain areas. To target neuroinflammation, we administered ABA to mice in the drinking water, for one month. We have previously published the beneficial effects of ABA as a safe anti-inflammatory treatment in rodent models of neurological alterations associated with metabolic syndrome [56,57] and Alzheimer’s disease mutations [46]. In addition, in this study, we compared the effect of the dopaminergic lesion and ABA administration between female and male subjects.

We found that dopaminergic lesion increased spontaneous activity in two-month-old males in agreement with findings in adolescent male mice [44,58], and in the DAT-KO rats (another model of dopaminergic dysfunction) [59]. Interestingly, the same lesion did not affect female’s activity, to our knowledge, this is the first comparative study, and our finding suggests a sex-dependent effect that may reflect the symptomatology in humans, where hyperactivity is found more prevalent in boys than girls [60]. Curiously, one month of ABA treatment increased spontaneous activity in lesioned females, whereas in males had the opposite effect, preventing the lesion-induced hyperactivity. Our results in males are in accordance with a recent study with hypertensive rats (considered a spontaneous ADHD model with regards to certain symptoms) that showed that curcumin could rescue hyperactive status in males [61]. Thus, our result suggests a different effect of the dopaminergic lesion in both sexes and a different response to ABA treatment. Further comparative studies are warranted to understand these sex-dependent differences and urge a revision of treatment for girls and boys, further supporting the need for precision medicine.

Other core symptoms, such as impulsivity and attention deficit were not evaluated in this study, further studies are necessary to understand the inflammatory etiology of diverse symptoms. In this study, we focused on related symptoms.

Exploration in mice is a marker of healthy development. We showed that in both females and males, 6-OHDA reduced the recognition memory, like in a previous report in one-month male mice [44]. However, one month of ABA treatment was not able to prevent this impairment. We hypothesised that longer treatments may be required, as we have previously observed in ABA intervention [46]. Spontaneous delayed alternation, a measure of spatial working memory, was similar in all groups. Previous reports have validated the impaired executive function in the 6-OHDA-induced lesion model by latent inhibition, and lower attention in the 5-choice serial reaction [58]. Our result indicates that dopaminergic lesion does not affect spatial working memory (exploratory behaviour and cognitive function), although affecting object recognition memory. This may be because spontaneous delayed alternation appears to rely on a variety of brain regions [62], that may have not been affected by the dopaminergic lesion.

In contrast to previous studies [44], we observed no differences in the anxiety levels of the 6-OHDA mice model, measured by the EPM. The differences can be attributed to the age difference in [28] and our study, they used 1- and we used 2 months-old mice. In humans, anxiety has been found to cooccur with ADHD core symptoms in approximately 50% of cases [63]. In addition, anxiety is highly dependent on the environment, so we cannot rule out external factors, even related to animal housing (i.e. our animals were habituated to experimenters and were housed in groups to reduce isolation stress, all factors that reduce anxiety).

In sociability tests, we observed that 6-OHDA females interacted more with co-specific than controls, whereas males had no difference with the lesion, suggesting an interaction between dopaminergic alterations of social functioning and sex. This feature has not been previously described since no female studies have been carried out in rodent models of ADHD. In humans, deficits in sociability have been associated with both genders with ADHD [64,65], whereas other studies report no differences[66]. Further studies are warranted to test sociability and social interactions in female and male animal models, to understand whether social deficits are a direct consequence of dopaminergic dysfunction or indirectly, a result of self-perceived stigma in humans.

Emotional dysregulation or sensory over-responsiveness is often found in ADHD children [67], [68]. These features can be used as predictors of ADHD and other psychiatric disorders [69]. Recent studies point to an association between pain sensitivity and emotional alterations in ADHD patients [70]. In our animal model, we have observed that female, but not male mice, had an increased pain sensitivity in thermal and mechanical stimulus, after the 6-OHDA lesion. Interestingly, studies on comorbidity between pain and ADHD and/or autism spectrum disorder, have reported altered pain sensitivity mostly in women [33]. Importantly, we did not find increased pain sensitivity in males, nor with thermal or mechanical stimulus, again pointing to a sex-dependent effect of brain dopaminergic depletion. Similarly, other studies in the male rat model reported no pain sensitivity to mechanical stimuli, but specifically to chemical stimuli [71]. However, other studies with the 6-OHDA lesion mice model of ADHD show increased sensitivity to mechanical and thermal stimuli [72]. Also, the Neurogranin knockout mice show hyperactivity and elevated pain sensitivity, together with behaviour traits associated with several human disorders [73]. In our study, control females showed a higher threshold to pain than males. Sex/gender differences in pain perception have been long observed in humans (reviews [74,75]), however, in rodent models, sex differences to different noxious stimuli vary greatly in different strains of rats and/or mice [76], suggesting an interaction between strain and environment, rather than sex and pain. There is evidence of an association between higher pain sensitivity and ADHD in boys [77], and adults with ADHD [35,36,78], however, other studies reported the opposite [79].

In our study, control females showed a higher threshold to pain than males. Sex/gender differences in pain perception have been long observed in humans (reviews [74,75]), however, in rodent models, sex differences to different noxious stimuli vary greatly in different strains of rat and/or mice [76], suggesting an interaction between strain and environment, rather than sex and pain. There is evidence of an association between higher pain sensitivity and ADHD in boys [77], and adults with ADHD [35,36,78], however, other studies reported the opposite [79].

Importantly, in females, one month of ABA treatment **prevented the pain sensitivity induced by the 6-OHDA lesion**. Moreover, ABA treatment promotes higher thresholds in males, implying a beneficial effect of ABA on pain sensitivity. This is further supported by recent reports showing that ABA ameliorates neuropathic pain, via reducing spinal cord inflammation [80].

To evaluate whether brain inflammatory status was associated with increased pain sensitivity, we next carried out the morphologic characterization of microglia in the anterior cingulate cortex (ACC) and the posterior insular cortex (pIC), as well as being components of the mesolimbic reward system, these areas are strongly implicated in pain processing [81,82]. Interestingly, females and males showed very different responses. We found that 6-OHDA injected females had proinflammatory microglia in ACC and pIC, and that one-month ABA treatment reduced this inflammatory phenotype in both areas, correlating to pain. Our result would agree with the notion that the ACC and pIC are primary targets for neuroinflammation. ACC has been associated with mood disorders and pain sensitivity [83]. Moreover, drug inhibition of pIC glia alleviates chronic pain in a rat model of nerve injury [84].

However, lesioned male mice displayed proinflammatory microglia in ACC, but not in pIC. Since males did not show alterations in thermal or mechanical stimulus sensitivity, this result may suggest that inflammation in pIC would correlate better with the behaviour observed. Interestingly, one month of ABA treatment increased the stimulus threshold in males but did not alter inflammatory microglia, suggesting that ABA may exert this effect via other areas and /or other mechanisms. Overall, our results indicate that dopaminergic reduction with 6-OHDA induces proinflammatory microglia in brain areas. This result agrees with the evidence suggesting that high dopamine levels induce antiinflammatory effects in microglia (via the activation of low-affinity dopamine receptors); while low dopamine levels can trigger inflammation (selectively stimulating the high-affinity dopamine receptors) [54]. Interestingly, we confirmed that a one-month treatment of ABA has an anti-inflammatory effect, but only on females, suggesting an important sex-dependent response to ABA. It remains to be determined if longer treatments would also improve microglia inflammatory status in males, as we have observed in previous studies [46].

Oxidative stress and neuroinflammation are also closely related to ADHD [85]. Apurinic/Apyrimidinic endonuclease/redox effector-1 (Ape1) is a vital mediator in redox signaling [86] in addition to its DNA repair function. Importantly, it is considered a hub for inflammatory-oxidative connections [87]. Thus, targeting APE1 has been considered a therapeutic option, either by reducing its function [88] or by restoring it [89,90]. This paradoxical effect of APE1 may be due to its pleiotropic effects, the toxic stimulus, or the cell type. It could also be related to the window of time, given the fact that APE1 is engaged when oxidative stress occurs, however its long-term reduced levels may increase vulnerability (i.g. as found in animal models of senescence, where reduced Ape1 correlates to increased mitochondrial DNA damage [91].

To our knowledge, this is the first study evaluating oxidative stress in brain areas of a rodent model of dopaminergic dysfunction. We found that 6-OHDA lesion reduces Ape1 expression in females’ ACC, and this reduction is rescued by one-month ABA administration. **Thus, our data would support that, in females, the increased pain sensitivity is associated with higher inflammatory and lower APE1 levels in ACC**. Other models of pain show that Ape1 levels are reduced in the spinal cord models of inflammatory pain [49]. Interestingly, in female pIC as well as males ACC and pIC, the lesion did not alter Ape11 intensity, but one-month ABA treatment increased significantly Ape1 intensity only in Sham animals, but not in 6-OHDA lesion. The effect in control mice would be in agreement with studies where Ape1 is upregulated by BDNF [92] since we have demonstrated that ABA administration can effectively increase BDNF expression [57]. Moreover, other studies have shown that resveratrol increased Ape1 levels in a rat model of aluminum-induced neuroinflammation [93]. Further studies are required to understand whether longer ABA treatment can rescue Ape1 levels in lesioned mice, and to understand the interaction between dopaminergic dysfunction and ABA action.

## 5. Conclusions

Our findings support a sex-dependent effect of 6-OHDA lesion, inducing higher pain sensitivity in females and increased spontaneous locomotor activity in males. In females, pain sensitivity was associated with inflammatory microglia and oxidative stress, in ACC and posterior IC. No inflammation was observed in these areas in males, suggesting an important sex-dependent interaction in the role of dopamine signaling in ACC-IC regulating the descending pain inhibitory pathway. Importantly, ABA treatment reduced inflammation and oxidative stress, alleviating pain hypersensitivity in females. Our data suggest a mechanism by which (some) ADHD patients may suffer from higher pain sensitivity, and points to higher risk for females. In males, ABA reduced hyperactivity, while not affecting neuroinflammatory status in these areas.

Our data strengthen the beneficial role of ABA and opens a potential alternative therapeutic intervention targeting inflammation for ADHD managment. Further studies are warranted to understand the sex differences in the responses to dopaminergic deficit and treatments.

## Author Contributions

Conceptualization, AMSP, NK, and ML; methodology, SSS, MMB; software, MMB, SSS; validation, MMB, SSS, AMSP; formal analysis, MMB, SSS, AMSP; investigation, MMB, SSS, AMSP; resources, AMSP; data curation, MMB, SSS, AMSP; writing—original draft preparation, AMSP; writing—review and editing, AMSP, SSS, MMB, NK, ML; visualization, AMSP; supervision, AMSP; project administration, AMSP; funding acquisition, AMSP.

All authors have read and agreed to the published version of the manuscript.

## Institutional Review Board Statement

The study was approved by the Ethics Committee of the University Jaume I. Protocol code 2020/VSC/PEA/0099); the date of approval 5/June/2020.

